# Temporal and cell-type specific SPAK-NKCC1 disruption following severe TBI in the developing gyrencephalic brain

**DOI:** 10.1101/2025.01.20.633889

**Authors:** Alexandra Hochstetler, Ya’el Courtney, Peace Oloko, Benjamin Baskin, Andrew Ding-Su, Tawny Stinson, Declan McGuone, Robin Haynes, Maria K. Lehtinen, Beth Costine-Bartell

## Abstract

Traumatic brain injury (TBI) is a leading cause of morbidity and mortality in infants and toddlers, with limited treatment options and persistent neurological sequelae. We developed a multi-pathoanatomic lesion multi-insult (MuLMI) severe TBI model in piglets that replicates age-dependent damage patterns to the cortical ribbon observed in human patients with less injury in postnatal day (PND) 7 “infant” piglets and more extensive tissue damage in PND30 “toddler” piglets. Given that neuronal chloride homeostasis influences excitability, seizure susceptibility, and edema, we examined the developmental and injury-induced regulation of key cation-chloride cotransporters and modulators: NKCC1 (sodium-potassium-2-chloride cotransporter), KCC2 (potassium-chloride cotransporter), and the regulatory kinase SPAK, which are biomarkers of neuronal chloride concentrations. This study is the first to define the spatiotemporal expression and phosphorylation profiles of these proteins in the developing piglet brain. We found a perinatal shift in the ratio of KCC2:NKCC1 across the brain, driven primarily by protein abundance, rather than transcriptional levels. We hypothesized that toddler piglets would exhibit an increase in cortical NKCC1 and SPAK causing hyperexcitability and perhaps explaining their more severe, unilateral cortical damage. Severe TBI induced a transcriptional increase in *Slc12a2* and *Stk39,* and a decrease in *Slc12a5* in toddler piglets, but not infant piglets. We further found that infant piglets, not toddler piglets, upregulated SPAK and Tyrosine Receptor Kinase B (TRKB) protein in cortex after TBI, with minimal changes in NKCC1 and KCC2. However, phosphorylated NKCC1 (pNKCC1) was significantly upregulated in surviving cortical neurons after TBI in infant piglets and was unchanged in toddlers, despite more severe injury. These findings suggest that cortical neuronal NKCC1 activation may play a role in post-traumatic excitability or resilience in the immature brain and identify NKCC1 and/or SPAK as a potential therapeutic target. In human tissue, the KCC2:NKCC1 ratio also increased postnatally, and TBI caused region and cell-type specific dysregulation of pNKCC1. Our results establish piglets as a valuable model for investigating age-specific mechanisms of pediatric TBI and for testing targeted interventions, particularly for infant populations where seizure control remains a major clinical challenge.

## INTRODUCTION

In infants and toddlers, most severe traumatic brain injury (TBI) results from abusive head trauma and is associated with high mortality and long-term neurological comorbidities [1]. Despite its prevalence, pediatric TBI remains understudied, and therapeutic options are limited. Young children of different ages exhibit distinct structural, biomechanical, and physiological vulnerabilities that shape injury severity and outcomes. Animal models must scale injury appropriately to brain volume, which increases significantly across postnatal development, especially in species like pigs and humans that undergo a perinatal brain growth spurt. Piglets are a high-fidelity model for pediatric TBI due to their gyrencephalic brain anatomy, developmental trajectory, and capacity to undergo intensive care interventions [1-5].

We employ a multi-pathoanatomic lesion, multi-insult TBI model (MuLMI) in piglets incorporating cortical impact, mass effect, subdural/subarachnoid hemorrhage, traumatic seizures, and transient apnea and hypoventilation under non-GABA-acting sedation to study the pathophysiology of severe TBI in young children. One example of MuLMI is the “hemispheric hypodensity” (HH) injury pattern observed in pediatric TBI patients: patchy, bilateral HH in infants and more unilateral HH in toddlers [6-8].

A critical contributor to these age-dependent injury patterns may be the “GABA switch”—the developmental transition in GABAergic signaling from excitatory to inhibitory, in part, driven by shifting expression and phosphorylation of two cation-chloride cotransporters: NKCC1 and KCC2. NKCC1 maintains high intracellular chloride ([Cl⁻]i), promoting depolarizing GABA responses, whereas KCC2 lowers [Cl⁻]i, enabling inhibitory GABA action [9, 10]. TBI may disrupt chloride homeostasis and exacerbate excitotoxicity [11-13]. Since standard seizure management for pediatric TBI often relies on GABA agonists [14, 15], an immature or disrupted chloride gradient may render these treatments ineffective or even harmful.

The timing of the GABA switch varies widely across species. In precocial species (e.g., sheep, guinea pigs), it occurs before birth; in altricial species (e.g., rodents), after birth; and in humans and swine, likely during the perinatal period [16-21]. Pinpointing this transition is essential for evaluating the translational validity of piglet models and for identifying vulnerable windows in pediatric brain injury. This is particularly relevant in diagnoses where seizure plays a pathogenic role, such as sudden unexpected death in childhood and early-life encephalopathies [22-26].

In this study, we defined the spatiotemporal expression of NKCC1 and KCC2 in the human and piglet brain and investigated their regulation following severe TBI. We found that TBI induces upregulation of phosphorylated NKCC1 (pNKCC1) in neurons of infant but not toddler piglets, with similar patterns observed in human tissue. Notably, neuronal pNKCC1 expression correlated positively with seizure duration, subarachnoid hemorrhage, and the extent of tissue damage: key features of our severe TBI model. These findings suggest that aberrant NKCC1 activity contributes to excitability and injury in early development, and that NKCC1 inhibition may represent a promising therapeutic target for mitigating damage after infant TBI. This work provides a valuable platform for understanding age-dependent mechanisms of pediatric brain injury and evaluating novel interventions in a translationally relevant large-animal model.

## MATERIALS AND METHODS

### Sample procurement

#### Animal procedures

All protocols and procedures performed were in accordance with the guidelines of the American Veterinary Association, the National Institutes of Health, and the United States Department of Agriculture. They were approved by the Institutional Animal Care and Use Committee at Massachusetts General Hospital.

#### Naïve swine tissue collection

##### Prenatal collections

Preterm piglets were collected from euthanized Yorkshire sows at approximately 30% gestation (ED36), 75% gestation (ED90), and 90% gestation (ED105). Pig fetuses were surgically removed from the amniotic sac following maternal euthanasia, rinsed under cold running water, brains were dissected on ice, and tissues were immediately flash frozen. In all cases, the time from euthanasia to sample collection was less than 4 hours.

##### Postnatal collections

7-day-old (PND7, “infant”) and 30-day-old (PND30, “toddler”) male Yorkshire piglets were selected for naïve tissue collection. Surgery and anesthesia protocols were employed as previously described (Costine-Bartell et al., 2019), adjusted to the age of the piglets. In brief, piglets were sedated with ketamine/xylazine/atropine and anesthetized by 1-1.5% isoflurane with medical oxygen. Piglets were intubated and mechanically ventilated with room air. An ear intravenous or external jugular vein catheter was placed to access venous blood and deliver saline infusion. In all cases, mean arterial blood pressure, oxygen saturation, heart rate and end-tidal CO2, respiratory rate, and core body temperature were managed for the duration of the procedures. A craniotomy was performed, and the dura was removed. Euthasol was delivered intravenously, and then fresh tissue dissection was performed immediately on wet ice. Samples (n=4/age) were collected from August 2023 to October 2023 and flash-frozen on dry ice.

##### Multi-focal severe TBI model injuries

In brief, a series of PND7 and PND30 male Yorkshire piglets from September 2016 to October 2019 were exposed to surgery and anesthesia protocols as previously described [6, 27]. A total of 23 piglets were used for fixed tissue and a total of 14 for fresh tissue. Immediately prior to the injury, piglets were switched to a seizure-permissive anesthesia protocol and subjected to a combination of injuries scaled to brain volume as previously described, including cortical impact, mass effect, subdural hematoma/subarachnoid hemorrhage supplemented with kainic acid, one-minute of apnea, and 10 minutes of hypoventilation. In the sham piglets, only the burr holes and duration of anesthesia were modeled. Piglets were managed in an intensive care unit for 24 hours after injury, where video EEG was recorded, beginning with scalp EEGs and then eventually with two 4-electrode epidural EEG strips (1 per hemisphere), resulting in a 4-channel bipolar montage as previously described [27].

Seizure length, subdural hematoma area, subarachnoid hemorrhage area, and tissue damage were analyzed as previously described [27]. Briefly, sections from FFPE blocks in 5 mm increments rostral to caudal in each hemisphere were analyzed for damage that included areas of “red neurons,” areas adjacent to red neurons with vacuolization of neuropil and blood vessels, and areas with acute hemorrhage. “Red neurons” were classified by the presence of all three of the following features: (1) cell shrinkage with perineuronal vacuolization, (2) cytoplasmic hypereosinophilia, and (3) nuclear pyknosis. This pattern of damage is termed “hypoxic-ischemic type injury,” though a myriad of pathways can lead to this common end-stage pattern of apoptosis/necrosis. Vacuolization alone was not included in the calculation of brain damage areas. Areas of damage were found to mostly overlap with albumin extravasation, expression of amyloid precursor protein, and matrix metalloproteinase-9, demonstrating vasogenic edema that extends far beyond the area of cortical impact [27, 28] where tissue edema might eventually lead to capillary and blood flow collapse. The percent of the hemisphere damaged presented here is a subset of previously published cohorts for the purpose of displaying the model and damage in the subjects used here for analyses of chloride salts membrane transporters.

#### Swine fresh tissue collection

Samples were requisitioned as freezer aliquots from prior studies (both sham and injured piglets) but are distinct aliquots from the same piglets published in [28]. For fresh frozen tissue, Euthasol was delivered intravenously, and then fresh tissue dissection followed immediately on wet ice, and samples were flash-frozen on dry ice. For perfused-fixed tissue, piglets were transcardially perfused with room temperature normal saline at a rate of ∼1L/min until complete surface vessel clearing followed by room temperature 10% neutral buffered formalin at the same rate until total body stiffness. Brains were obtained and post-fixed in 10% neutral buffered formalin for 3-5 days until appropriate tissue stiffness was achieved.

#### Human tissue collection

Lysates from frozen hippocampus at the level of the lateral geniculate nucleus were collected from either the NIH NeuroBioBank or the San Diego County Medical Examiner’s office, the latter available for research under the auspice of the California Code, Section 27491.41. All infants died suddenly and unexpectedly with the cause of death determined at autopsy and adjudicated within the laboratory for research purposes (**Table 1**). Unstained tissue sections from FFPE blocks were procured from autopsy brain samples (TBI or non-TBI control) performed at the Neuropathology Department at the University of Edinburgh. Permission to use the brain tissue was given according to the guidelines required by the respective ethics committees of these institutions: the Boston Children’s Hospital and Massachusetts General Hospital Institutional Review Boards. Demographic information of all human samples is detailed in **Table 1**.

**Table 1:** Human Sample Demographics.

|  | Sample IDs | PNA (GA) | Sex | Diagnosis | PMI (hr) |
| --- | --- | --- | --- | --- | --- |
| **Frozen/FFPE Hippocampus (Figure 2)** | 1 | 0.4 (40) | Female | Fatty acid oxidation disorder | 15 |
|  | 2 | 0 (41) | Female | Neonatal bradycardia with arrest and meconium aspiration | 12.8 |
|  | 3 | 17 (26.5) | Female | Acute asphyxiation | 9 |
|  | 4 | 12 (32) | Male | Acute asphyxiation | 9 |
|  | 5 | 5 (40) | Male | Acute asphyxiation | 5 |
|  | 6 | 11 (36) | Female | Acute asphyxiation | 27 |
|  | 7 | 13.5 (37.5) | Female | Acute asphyxiation | 29 |
|  | 8 | 16 (40) | Female | Acute asphyxiation | 9 |
|  | 9 | 26 (36) | Male | Acute asphyxiation | 9 |
|  | 10 | 33.4 (38.5) | Male | Intussuseption of inverted Merkel's diverticulum | 13 |
|  | 11 | 35 (39) | Female | Acute asphyxiation | 29 |
| **FFPE Non-TBI Control (Figure 6, Supplemental File 1)** | Non-TBI-1 | 0 (23) | Unknown | Normal brain development; labor/pregnancy complications | <24 |
|  | Non-TBI-2 | 0 (27) |  |  |  |
|  | Non-TBI-3 | 0 (39) |  |  |  |
|  | Non-TBI-4 | 0 (40) |  |  |  |
|  | Non-TBI-5 | 4 month | Male | Sudden unexpected death in infancy | 24 |
|  | Non-TBI-6 | 5 month | Male | Cystic fibrosis | 3.33 |
|  | Non-TBI-7 | 4 year | Female | Acute peritonitis | 24 |
| **FFPE TBI (Figure 6, Supplemental File 1)** | TBI - 1 | 3 month | Female | Blunt force head injury | 23 |
|  | TBI - 2 | 3 year | Female | Traumatic axonal injury, subdural hematoma | 24 |
|  | TBI - 3 | 3 year | Male | Road traffic accident, contusions, cerebral edema | 24 |
|  | TBI - 4 | 9 year | Female | Road traffic accident, contusions, cerebral edema, subdural hematoma, global ischemia | 24 |
|  | TBI - 5 | 9 year | Male | Road traffic accident, contusions, cerebral edema, subdural hematoma, global ischemia | 24 |
|  | TBI - 6 | 10 year | Female | Road traffic accident, contusions, cerebral edema | 72 |

### Standard laboratory techniques

#### Western Blotting

Fresh frozen samples were manually homogenized in N-PER lysis buffer (ThermoScientific #87792) supplemented with 1X Halt (ThermoScientific #78440) protease and phosphatase inhibitor cocktail and lysed at 4°C for 60 minutes with end-over-end agitation. Lysates were spun at 16,000 x g for 10 minutes to pellet debris. Supernatants were aliquoted and Pierce BCA assay was used protein concentration (ThermoScientific #23250). 20µg of the sample was reduced with 5X Laemmli Sample Buffer (BioLegend #426311) supplemented with B-mercaptoethanol (5%) and diluted with double distilled water to an equivalent volume. Samples were heated at 70°C for 5 minutes, loaded into 4-15% TGX Stain-Free gels (Biorad # #4568086) and separated at 180V for 45 minutes. Gels were transferred to 0.2 µm nitrocellulose membranes at 1.0A for 30 minutes using the Bio-Rad Trans-Blotter® system. Membranes were stained with Ponceau-S total protein stain blocked with EveryBlot blocking buffer with gentle agitation. Membranes were incubated with primary antibody diluted in EveryBlot blocking buffer washed with 1X Tris Buffered Saline-0.1% Tween-20 (TBST) and then incubated with secondary antibody diluted in EveryBlot blocking buffer. Membranes were washed with TBST and TBS, exposed with chemiluminescent reagents (Cytiva-Amersham #RPN2235), and visualized on a Bio-Rad Chemi-Doc Imager. For antibody validation, full complete membranes were first probed with each antibody to evaluate non-specific bands, specific bands and dimers, and to optimize dilutions. In the case of pNKCC1 and NKCC1 antibodies, our laboratory has performed in-house knock-down validation. All validation methods are presented in **Table 2**. For publication gels, technical duplicate membranes were cut at 80kDa to allow probing of 1) pSPAK, 2) SPAK, and 3) beta-actin (below 80kDa) simultaneously with 1) pNKCC1, 2) NKCC1, and 1) pKCC2, 2) KCC2 (above 80kDa). In Figure 1 and 6, band intensities were normalized to total protein loaded (Ponceau stain) for the graphs, since beta-actin and beta-tubulin were found to be poor loading controls across age and region. In Figures 3 and 5, band intensities were normalized to both beta actin and total protein, and beta actin normalization is shown in the figure. Membrane stripping was performed with ReStore™ Western Blot Stripping Solution and protocol (ThermoScientific #21059). All western blot data presented and quantified in this manuscript was the result of a minimum of three technical replications on different days.

**Figure 1:**
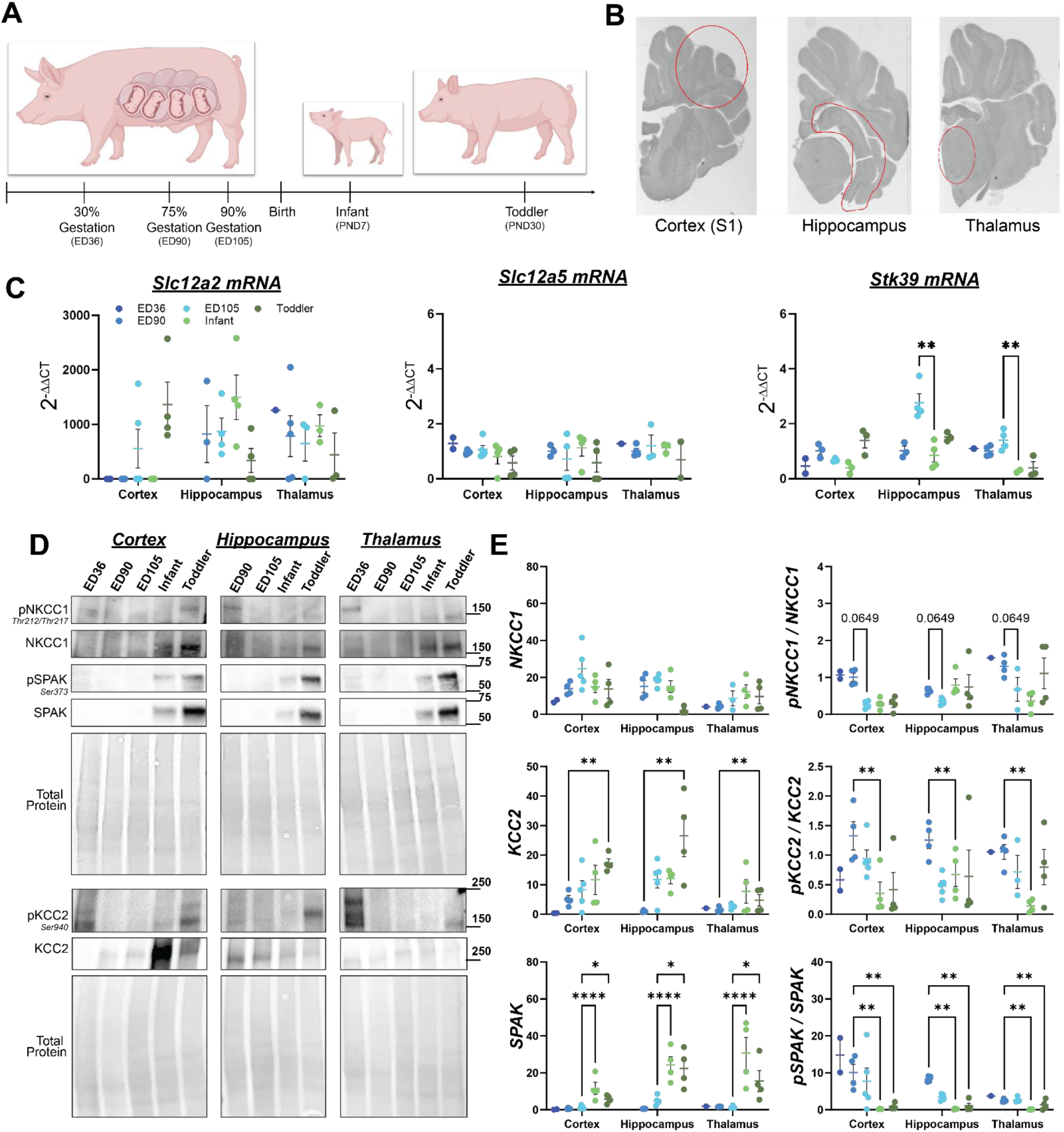
Proxy markers of transition to inhibitory GABA increase perinatally in swine, particularly across cortex. ***A***, schematic of samples represented across gestation and postnatal ages. ***B***, regions collected for analysis. ***C,*** quantitative PCR analysis of samples across ages (ED36, ED90, ED105, Infant, Toddler) and regions (cortex, hippocampus, thalamus) for *Slc12a2* (encoding NKCC1), *Slc12a5* (encoding KCC2), and *Stk39* (encoding SPAK). 2-way ANOVA for *Stk39* expression across ages and regions: main effect of age p=0.0007, main effect of region p=0.0008; post-hoc multiple t-test p<0.01. D, representative western blots for protein and phospho protein abundance, total protein stain as loading control. E, quantitation of western lots for samples across ages and regions for total protein (left) and ratio of phosphorylated to total protein (right). KCC2: 2-way ANOVA main effect of age p=0.0002, main effect of region p=0.0043; post-hoc multiple t-test **p<0.01. SPAK: 2-way ANOVA main effect of age p<0.0001, main effect of region p-0.008; post-hoc multiple t-test *p<0.05,****p<0.0001. pKCC2/KCC2: 2-way ANOVA main effect of age p=0.0015, main effect of region n.s.; post-hoc multiple t-test **p<0.01. pSPAK/SPAK: 2-way ANOVA main effect of age p<0.0001, main effect of region p=0.0104; post-hoc multiple t-test **p<0.01. pNKCC1/NKCC1: 2-way ANOVA main effect of age p=0.0012, main effect of region p=0.018; post-hoc multiple t-test p=0.0649.

**Figure 2:**
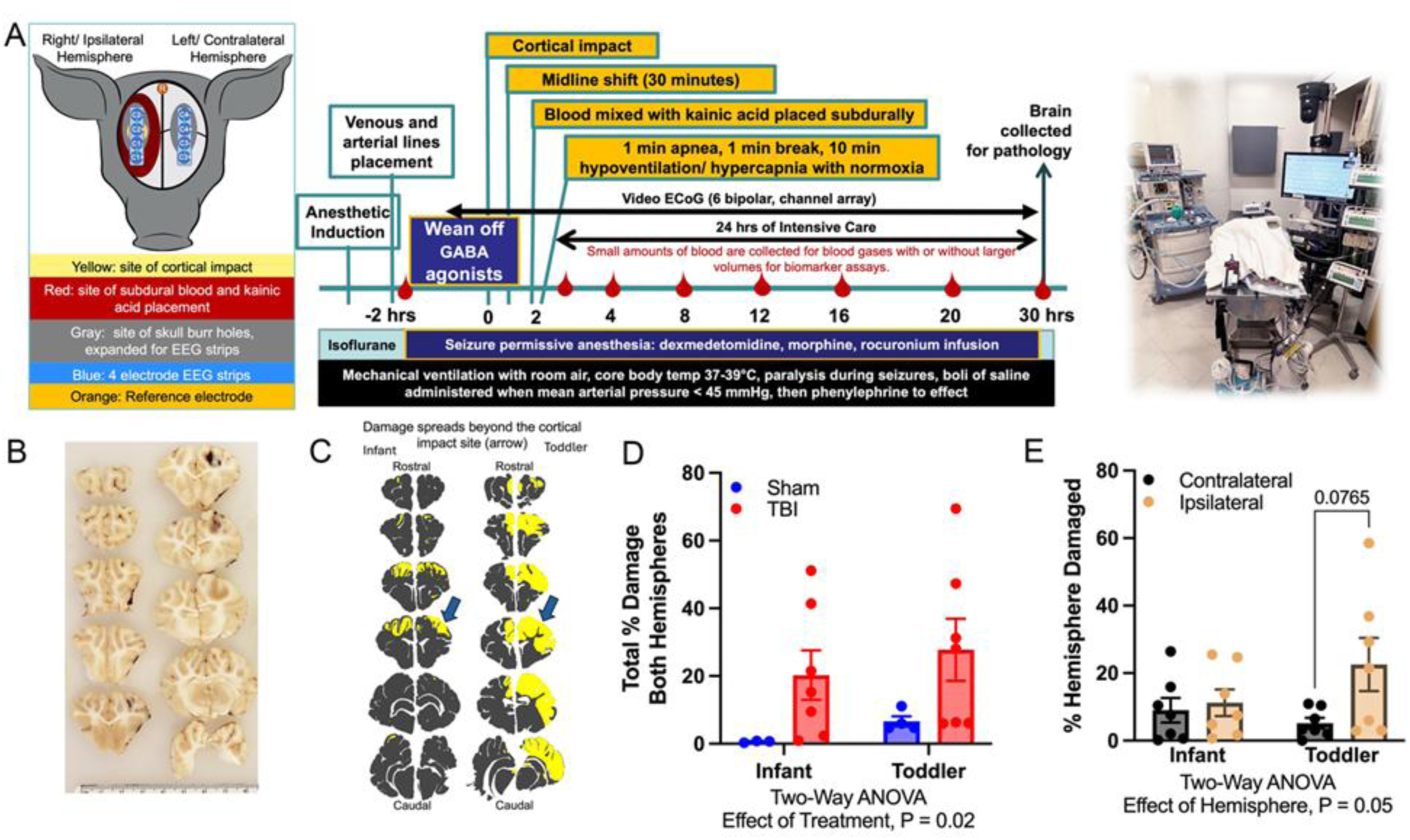
Multi-factorial, severe TBI piglet model injuries result in age-specific pathology patterns. ***A*,** Left:The array of injuries displaying the location of various injuries and insults per hemisphere and the electrode EEG strips. **Center:** The timeline of the injuries and insults performed after weaned from GABA agonists, along with intensive care, lasting approximately 24 hours to allow the injury to evolve. **Right:** Photograph of our ICU equipment where piglets are sedated, intubated, and ventilated. ***B*,** 5 mm brain slabs (right is right), and the subarachnoid hemorrhage can be seen along with dusky tissue and a swollen right hemisphere. ***C*,** Maps displaying the damaged areas (yellow) rostral to caudal and site of cortical impact (arrow). Damage spreads beyond the site of the cortical impact as we now hypothesize that seizures and subarachnoid hemorrhage synergistically drive tissue damage. ***D*,** In the cohort studied here, the amount of damage was greater in piglets receiving severe TBI injuries in both infants and toddlers (p=0.02). ***E,*** The percentage of the hemisphere damaged (microscopically determined in one section per 5 mm slab and mapped) exhibited age-dependent patterns where the ipsilateral hemisphere had greater damage than the contralateral hemisphere in toddlers (p=0.05).

**Figure 3:**
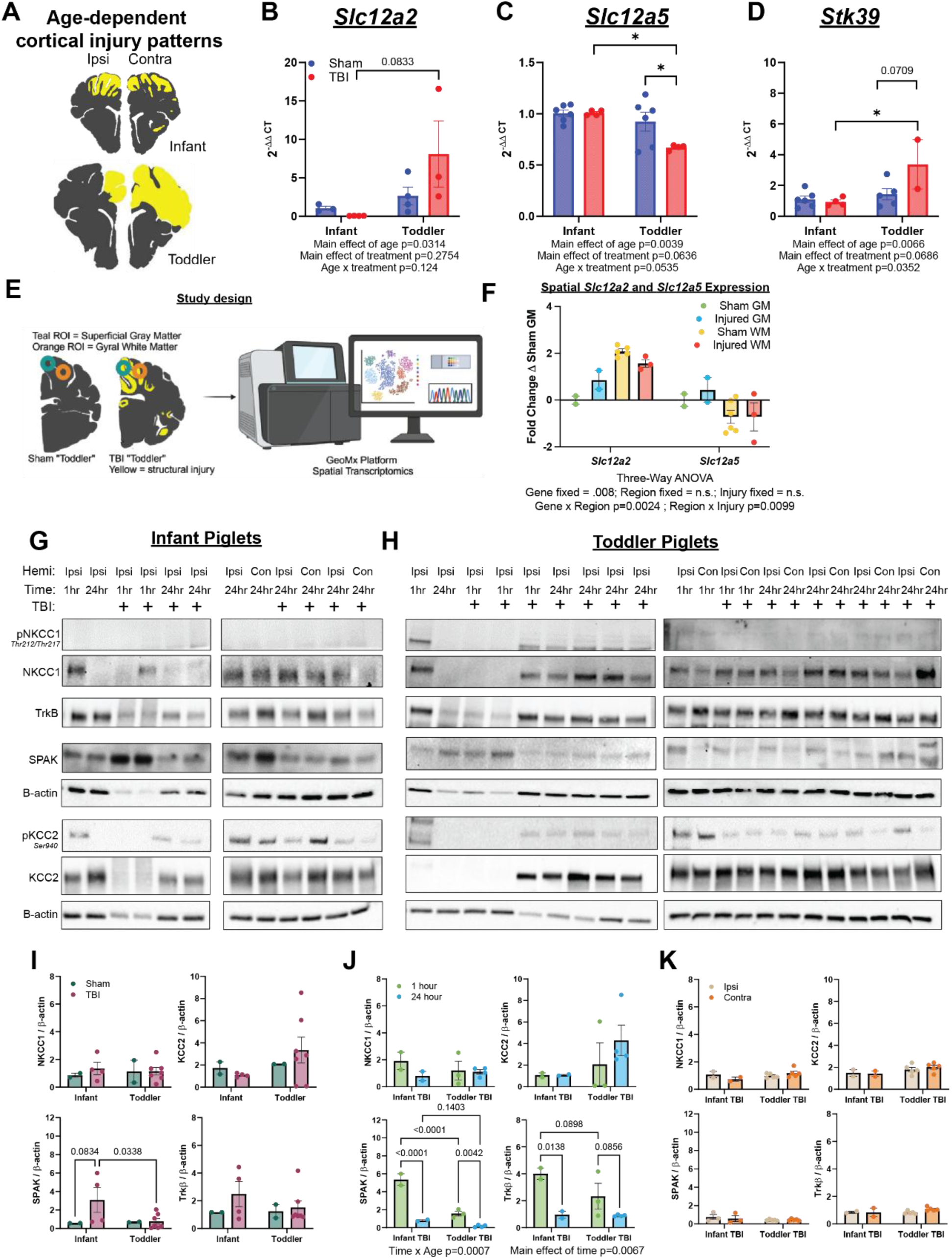
Severe TBI induces a reversion of markers of immature GABA in “infants” and upregulation of maturation in “toddlers”. Representative patterns of hypoxic-ischemic injury at the level of the somatosensory cortex between ages (***A***, yellow = injured tissue). Quantitative PCR analysis for transcripts *Slc12a2 (**B**)*, *Slc12a5 (**C**)*, and *Stk39 (**D**)* due to age and TBI injuries in piglets, presented as fold change from sham infant piglets. Two-way ANOVA results are below graph, and post-hoc multiple t-tests are included on graphs where applicable (*p<0.05, **p<0.01). ***E***, Study design and ROI selection for GeoMx Spatial Transcriptomics. ***F***, selected relative expression of *Slc12a2* and *Slc12a5* across regions from sham animals and animals with severely injured tissue. Three-Way ANOVA performed to analyze transcriptional levels across gene, region, and injury. Western blots from cortical lysates in infant (G) and toddler (H) piglets probed for pNKCC1, NKCC1, TrkB, SPAK, pKCC2, and KCC2, normalized to beta actin, quantified by band densiometry and graphed with regard to sham v. TBI (I), time for TBI injuries (J), and hemisphere for TBI injuries (K). Two-way ANOVA results are below graph, and post-hoc multiple t-tests are included on graphs where applicable with exact p values.

**Figure 4:**
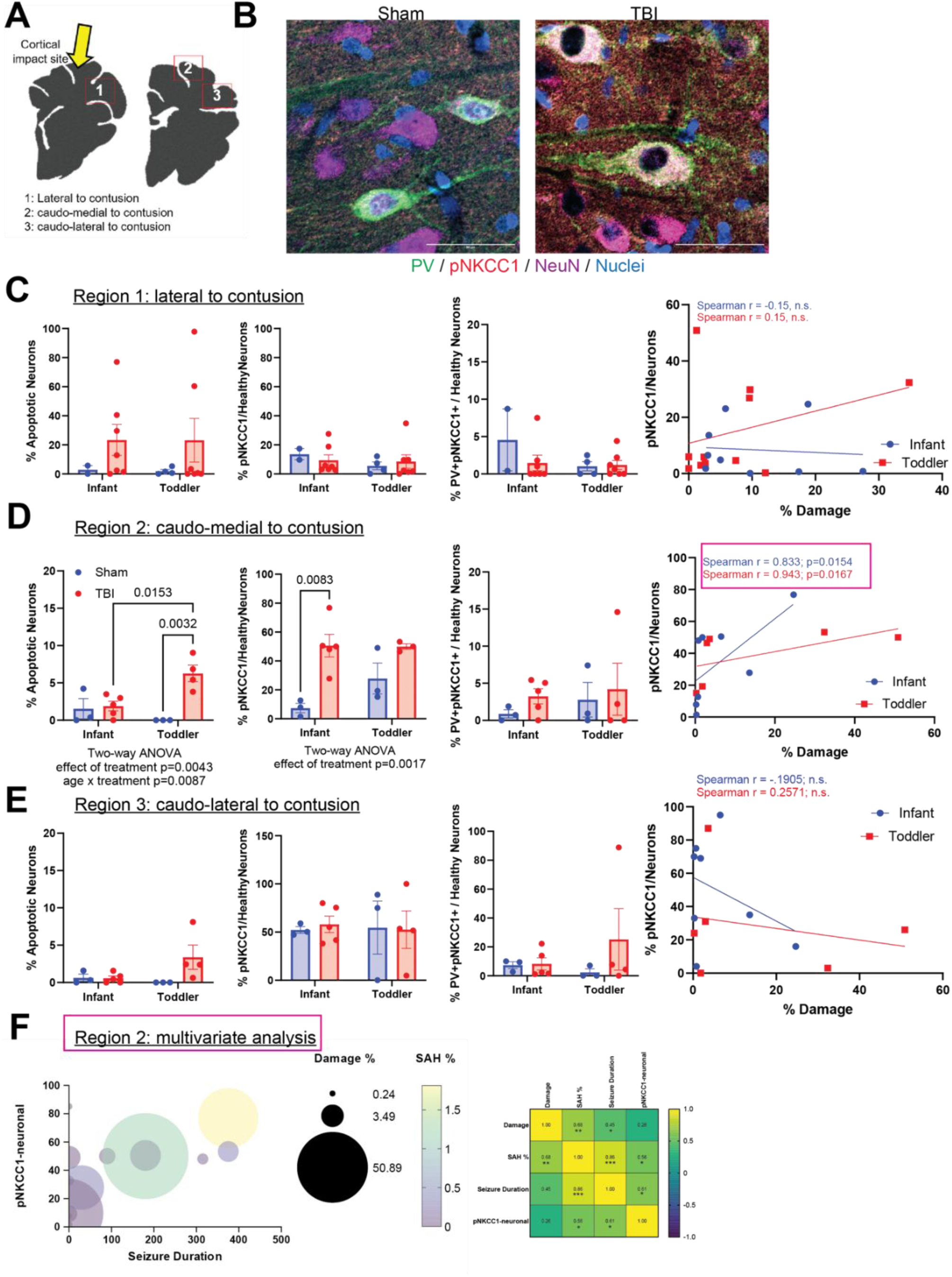
Cortical non-apoptotic, PV^-^ neurons upregulate pNKCC1 caudo-medial to contusion site in TBI-induced spreading hypoxic-ischemic injury and are associated with burden of tissue damage across ages and burden of subarachnoid hemorrhage and seizure duration in infant piglets with TBI. Schematic of ROIs for analysis in proximity to contusion site, *A*. Immunohistochemical staining for pNKCC1 (red), parvalbumin (green), NeuN (magenta), and nuclei (white), *B*. Cell counts expressed as percentages per ROI, with ROI 1 in *C*, ROI 2 in *D*, and ROI 3 in *E*. In all cases, the **left** dot plot presents % of neurons that were apoptotic, the **middle** dot plot presents the % of healthy neurons expressing pNKCC1, and the **right** dot plot presents % of healthy neurons co-expressing parvalbumin and pNKCC1. The scatter plot with spearman correlation on **far right** examines whether pNKCC1 neuronal expression correlates with damage. Two-way ANOVA with post-hoc multiple t tests employed for all comparisons and significant findings for main effect of age or treatment reported in graphs. Since a significant correlation with damage was found in region 2, *F* shows the multivariate analysis considering seizure duration, subarachnoid hemorrhage %, and damage % with Spearman correlation matrix presented and significant interactions marked with asterisk scheme, ** p<0.01, * p<0.05.

**Figure 5:**
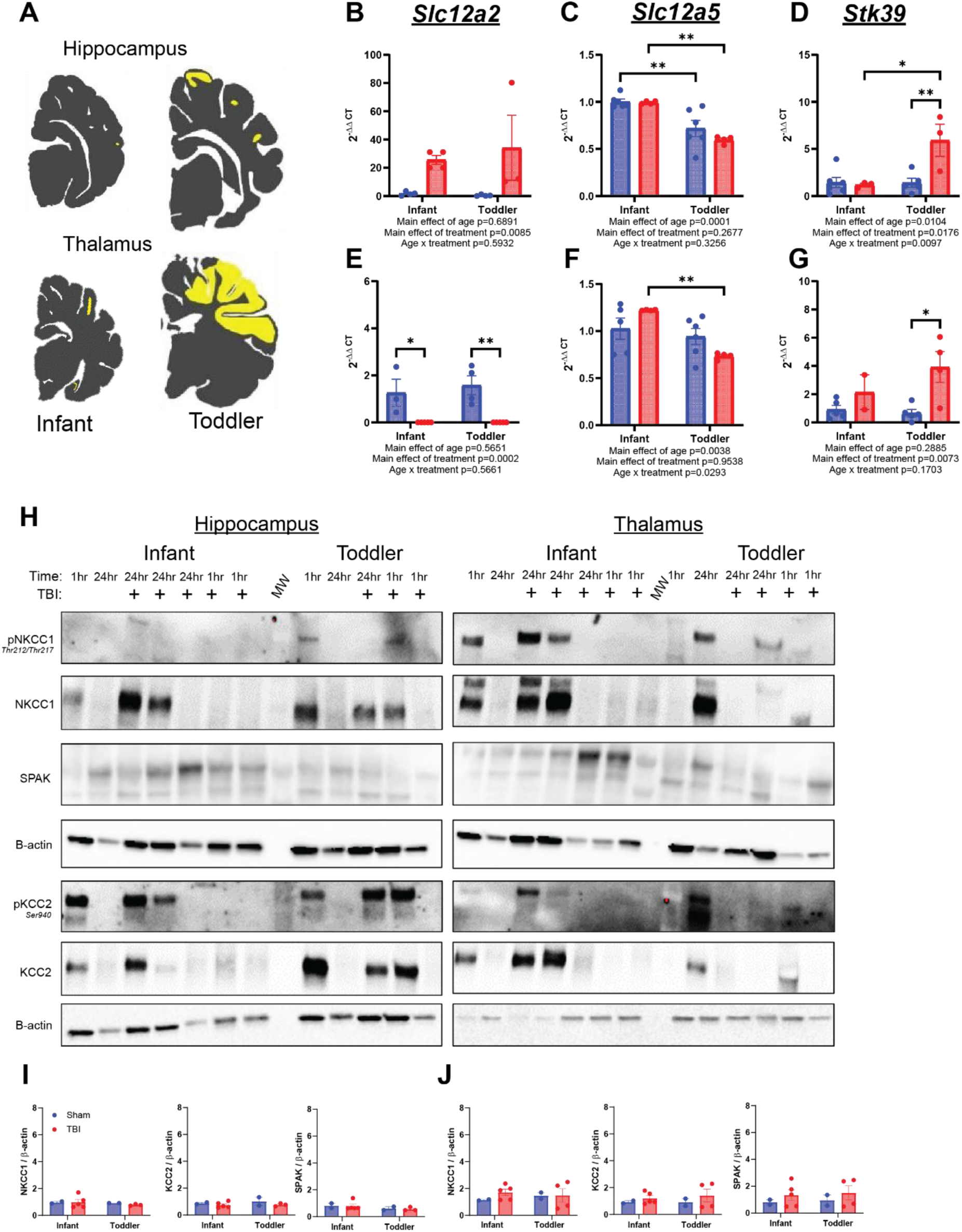
Protein expression of NKCC1, KCC2, and SPAK in the hippocampus and thalamus were minimally sensitive to severe TBI injuries in piglets despite persistent transcriptional changes. Representative patterns of minimal hypoxic-ischemic injury at the levels of the hippocampus and thalamus between ages (*A*, yellow = injured tissue). Quantitative PCR analysis for transcripts *Slc12a2 (B, E)*, *Slc12a5 (C, F)*, and *Stk39 (D, G)* due to age and TBI injuries in piglets, presented as fold change from sham infant piglets in hippocampus (*B-D*) and thalamus (*E-G*). Two-way ANOVA results are below graph, and post-hoc multiple t-tests are included on graphs where applicable (*p<0.05, **p<0.01). Western blots for all biological replicates for infant and toddler piglets receiving sham or severe TBI injuries in hippocampus and thalamus (*H*) and quantitated below (*I, J*).

**Figure 6:**
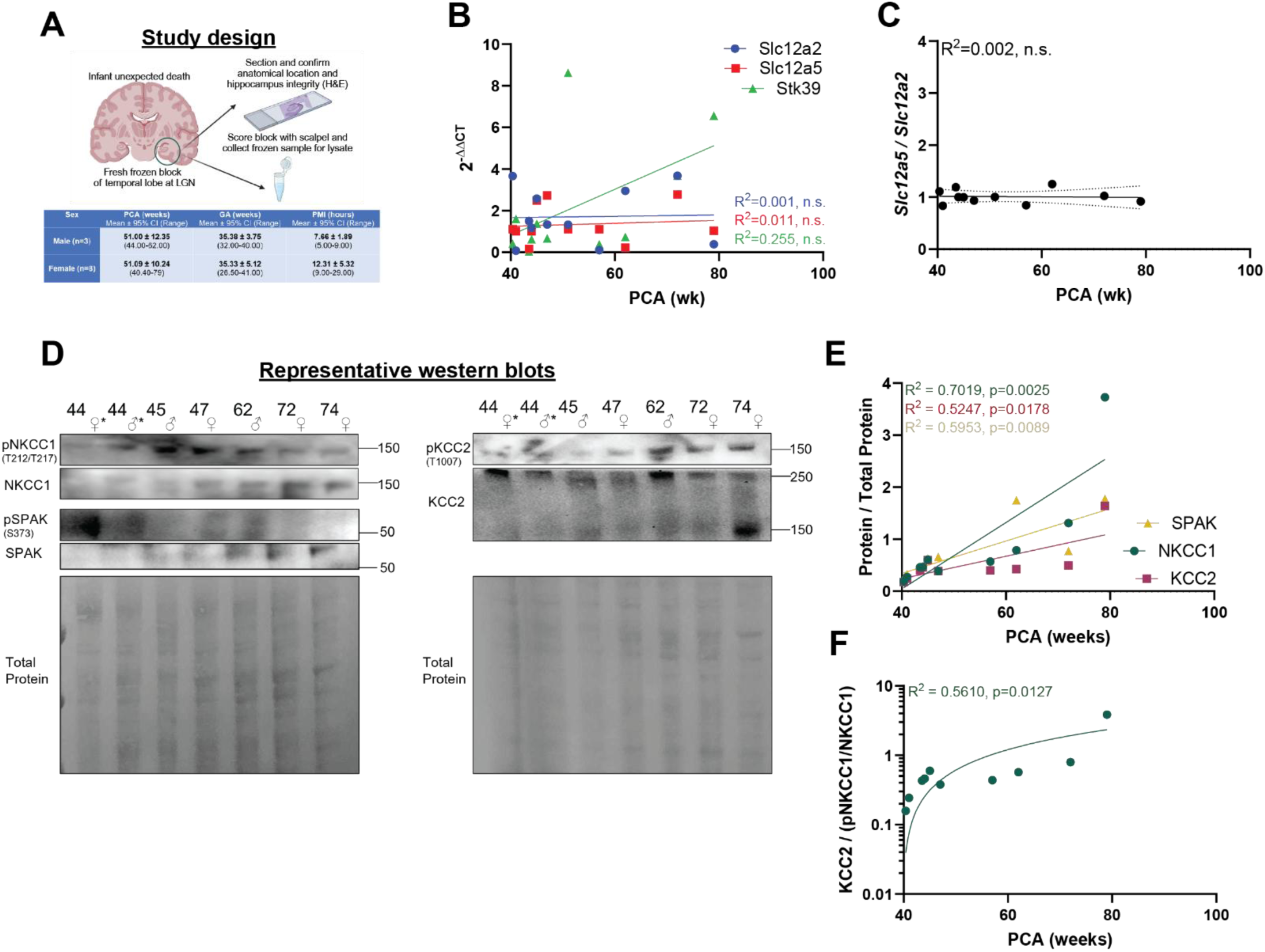
Postnatal GABAergic maturation occurs in human hippocampus. Schematic of study design, sample collection protocol, and sample demographics (***A***). Quantitative PCR analysis for transcripts *Slc12a2, Slc12a5,* and *Stk39 (**B**)* across postconceptional age (PCA) presented as fold change from youngest sample. Ratio of Slc12a5 to Slc12a2 as a function of PCA (***C***). Representative western blots of hippocampal lysates (***D***), quantified in (***E***) as a function of PCA. SPAK, NKCC1, and KCC2 all demonstrate positive linear correlations with PCA. Ratio of KCC2 : (pNKCC1/NKCC1) (***F***) also demonstrates a significant positive correlation with PCA. In all graphs, linear regression analysis was used. R2 values are shown on the graphs along with a p-value for whether the slope is non-zero.

**Table 2:** Antibody information.

| Antibody | Species | Target | Company | Catalog # | Validation Method | WB | IHC |
| --- | --- | --- | --- | --- | --- | --- | --- |
| NKCC1 | Rabbit | Synthetic Arg77 of mouse NKCC1 | Cell Signaling Technology | 14581 | KO lysate | 1:1000 |  |
| pNKCC1 | Rabbit | Human pNKCC1 Th212/217 | Millipore | ABS1004 | KO lysate | 1:500 | 1:250 |
| SPAK | Mouse | GST-tagged c-terminus of human STK39 | Millipore | MABS178 | (+) ctrl human brain tissue | 1:1000 |  |
| pSPAK | Rabbit | Human pSPAK Ser373 | Millipore | 07-2273 | Hypotonic HEK293 cells | 1:500 |  |
| KCC2 | Rabbit | Synthetic peptide | Abcam | ab259969 | (+) and (-) ctrl tissue lysates | 1:2000 |  |
| pKCC2 | Rabbit | Synthetic Ser940 | Invitrogen | PA5-95678 | KO lysate | 1:1000 |  |
| Parvalbumin | Sheep | E. coli-derived recombinant human Parvalbumin alpha Ser2-Ser110 | Invitrogen | PA5-47693 | (+) ctrl brain tissue |  | 1:100 |
| NeuN | Conj 647 | Synthetic peptide | Abcam | ab190565 | (+) ctrl brain tissue |  | 1:500 |

#### Quantitative, Real-Time PCR

Samples were processed for RNA extraction using the Total RNA Miniprep Kit (New England BioLabs #T2010), evaluated for concentration >50ng/µL and a quality ratio of A260/280 of >2.0. RNA was prepared for cDNA synthesis with +reverse transcriptase and –reverse transcriptase reactions using the LunaScript® RT Supermix Kit (New England Biolabs #E3010) using quality control metrics of concentration >100ng/µL and A260:280 ratio >1.8. Samples were diluted to equivalent cDNA loading and probed in triplicate with Taqman™ reagents for quantitative gene expression using fast run chemistry on an Applied Biosystems machine. In all cases, duplicate -RT and singlet no template controls were run for each sample. Probe information is detailed in **Table 3**.

**Table 3:** Taqman probe information.

| Target | Species | Sequence |
| --- | --- | --- |
| *Nkcc1* | *Human* | *Hs01074325_m1* |
| *Kcc2* | *Human* | *Hs00221168_m1* |
| *Stk39* | *Mouse* | *Mm01270753_m1* |
| *Reference Gene (18s)* | *N/A* | *Hs99999901_s1* |

#### Spatial Transcriptomics

Spatial transcriptomics was conducted on anatomically matched formalin-fixed, paraffin-embedded (FFPE) brain sections from a sham 30-day-old piglet and a 30-day-old piglet with severe traumatic brain injuries using NanoString GeoMx® Digital Spatial Profiling technology. Regions of interest (ROIs) were selected to capture key anatomical areas relevant to age-dependent TBI pathology and neuroinflammation. Following ROI selection, samples underwent GeoMx® platform analysis, where regions were scanned, and digital counts were obtained for gene targets of interest. Signal thresholding at a 3% cutoff was applied to exclude low-expression signals, ensuring that only significant transcript data were included in downstream analysis. Normalization was performed using internal controls (External RNA Control Consortium) provided by the GeoMx® platform to account for technical variation across slides and samples. Upper quartile normalization was performed after filtering the dataset to remove targets below the limit of quantitation.

#### Histology

Following immersion fixation, brains were washed with 1X phosphate-buffered saline (PBS), paraffin embedded and sectioned at 5-µm thickness and mounted onto silanized glass slides. Immunofluorescence was performed on these sections using antibodies detailed in **Table 2**. Briefly, sections were deparaffinized in xylenes, rehydrated, and then subjected to antigen retrieval in boiling 10mM citric acid-Tween pH 6.0 for 20 minutes. Staining protocols are presented in more detail below. In all cases, negative control sections were incubated in corresponding species-specific IgG at an equivalent concentration to the primary antibody dilution during the primary antibody incubation step. For figures where quantitation is presented, all slides evaluated were stained together in a single batch to minimize signal variability. Slides were imaged on a Zeiss Axio Observer D1 inverted microscope (fluorescent, 20x air objective) or a Zeiss LSM980 (20x air or 63x oil objective) at the Boston Children’s Hospital Cellular Imaging Core. Images were processed and assembled using ZEN Blue, ImageJ software, and Adobe Illustrator. For quantitation presented in **Figures 4**, **7, and S2**, the examiner performed cell counts on images that had been blinded and randomized.

**Figure 7:**
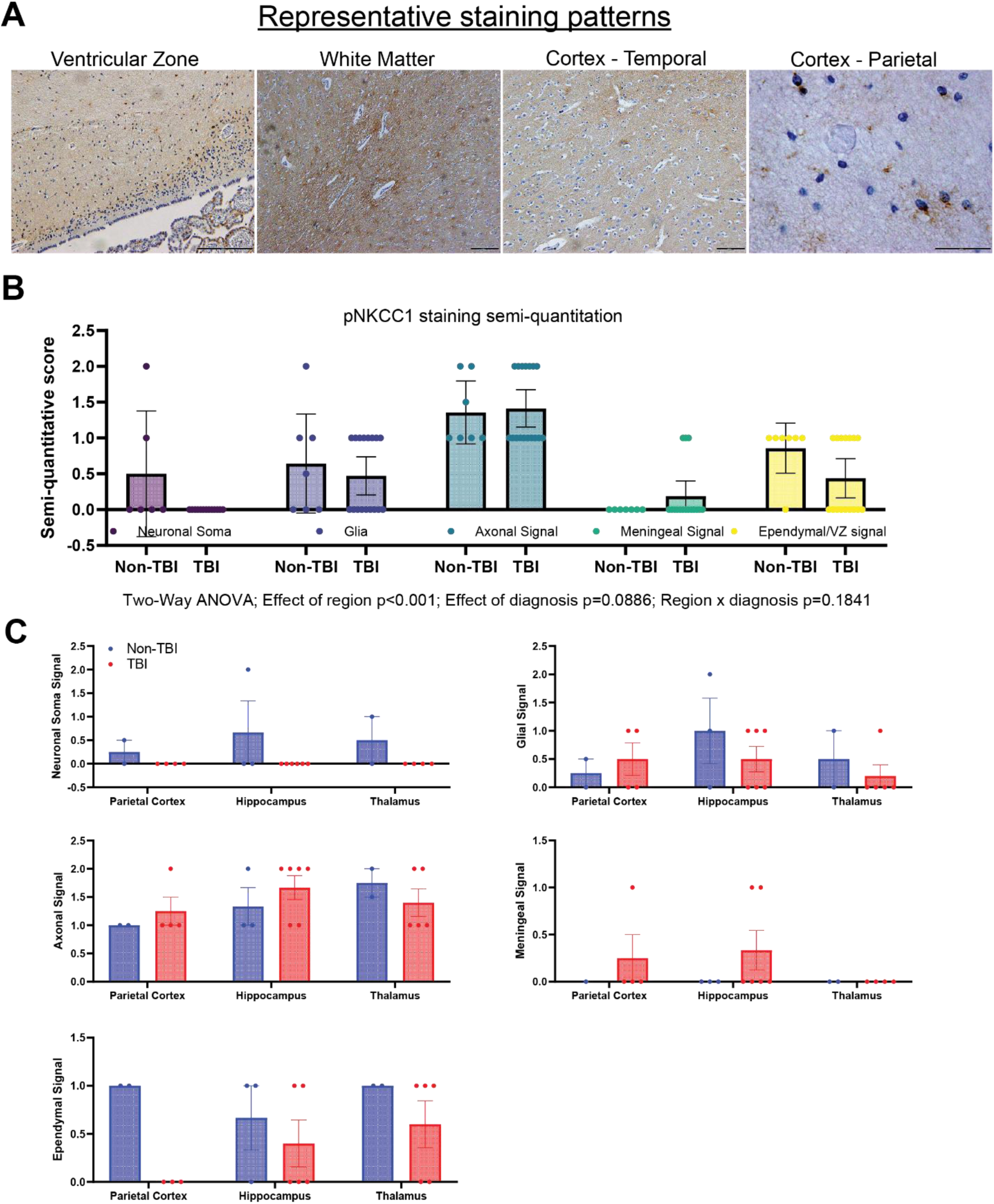
TBI induces heterogeneous cellular and regional changes to pNKCC1 expression and localization. Representative pNKCC1 staining patterns throughout human brain (***A***), semiquantitated as an effect of cell type (***B***) and cell type x region (***C***). Two-Way ANOVA in (***B***) with effects listed below graph.

##### Chromogenic Immunohistochemistry

For chromogenic stains, a peroxidase quenching step was performed immediately prior to antigen retrieval. Sections were then permeabilized for 5 minutes with 1% PBS-Tween-20 and blocked for 1 hour using normal horse serum diluted in Super Sensitive Wash Buffer (BioGenex, HK583-5KEGP). Sections were incubated in primary antibodies diluted in blocking buffer for 2 hours at room temperature. Slides were washed with 1% PBS-Tween-20 and then incubated in the secondary antibody diluted in blocking buffer for 30 minutes. Sections were washed and subjected to antibody visualization with Vectastain® ABC and DAB reagents (VectorLabs), counter stained with Mayer’s hematoxylin plus bluing reagent, and then dehydrated, cleared, and mounted with resin-based medium (Cytoseal™ 60 Mounting Medium, Epredia). Slides were sent coded to a blinded pediatric neuropathologist (DM) for analysis.

##### Fluorescent Immunohistochemistry

Sections were permeabilized for at least 10 minutes with 0.3% Triton X-100 in PBS and subsequently blocked for at least 40 minutes at room temperature with blocking solution (3% donkey serum and 2% BSA). Following, sections were washed with PBS and incubated in primary antibody diluted in blocking solution for 2 hours at room temperature. Next, sections were washed again and incubated in secondary antibody diluted in blocking solution for 1 hour at room temperature. Sections were washed and exposed to Hoechst 33342 nuclear stain for five minutes and mounted with an aqueous mounting medium (Fluoromount-G™).

#### Study design and rigor

The evaluation of the GABA switch in the naïve swine brain was planned prospectively, and the maximum biological replicates were collected as allowed by protocol restrictions. Postnatal samples were analyzed first to determine if the study was sufficiently powered. Following this power analysis, all naïve swine samples were analyzed together to generate bar plots as presented. The effect of TBI injuries in swine on the cation chloride cotransporters was characterized using aliquots and sections from a prior cohort where the main findings of age, injury, seizure medications, and mechanisms of vasogenic edema had been published. The original cohorts were randomized to treatment and exclusion criteria as previously published. Real-time PCR was performed in triplicate with the appropriate controls described above. Western blots were validated by running full length blots with the appropriate controls described above for antibody specificity and were run in technical duplicate (minimum of three times on two duplicate gels per run). For the spatial transcriptomics experiment, Figure 3, the region of interest was the experimental unit. Cell counts from Figures 4, 7, and S2 were performed by counters blinded to treatment as described above, with auditing on every 10^th^ section. Counters were within 10% of each other. The evaluation of frozen human brain sections was performed in the BCH Department of Pathology. A series of infants who died from non-TBI causes was used. Samples with an extended postmortem interval (PMI) (>30 hours) were excluded because prolonged PMI positively correlated with NKCC1 expression, likely a reflection of tissue autolysis in our pilot analysis. The effect of TBI injuries in humans on cation chloride cotransporters was examined using autopsy samples (**Table 1**) and semi-quantitatively evaluated by a board-certified forensic neuropathologist (DM) blinded to cause of death and age.

### Statistical analysis

Graphing and statistical analyses were completed in GraphPad Prism. Parametric statistical tests two- or three-way ANOVA considering age, treatment, co-variates, and/or region as applicable and described in the figure legends. Single variate analysis utilized Pearson correlation and linear fit (R^2^) and/or p values are displayed on the graph where applicable. Multivariate analysis was performed when applicable and multiple linear regressions correlation matrices are displayed on the figure. Exact p values are included on the graphs. For non-parametric data, Spearman correlation, Wilcoxson Signed Rank Test or Kruskal-Wallis Test by Ranks considering age, treatment, and/or region as applicable were performed as described in the figure legends.

## RESULTS

### Establishing the swine “GABA switch” with high spatial and temporal resolution

We examined the expression and abundance of key chloride cotransporters (*Slc12a2*: NKCC1 and *Slc12a5*: KCC2) and a regulatory kinase (*Stk39:* SPAK) in the American Yorkshire pig (*Sus scrofa domesticus*) across several developmental stages and brain regions, aiming to map how these changes might align with a prenatal, perinatal, or postnatal GABA switch (**Figure 1A, B**). We used whole tissue lysates of specific brain regions (**Figure 1C**) and applied quantitative PCR of tissue lysates, and western blot of total protein and phospho-protein (**Figure 1D, E**) to assess this question.

#### mRNA expression of Slc12a2, Slc12a5, and Stk39 across development and region in swine

In many studies, the relative mRNA expression of these transporters and a ratio of *Slc12a5: Scl12a2* have been utilized as a marker for the “GABA switch” (ex. [17]). Our intent was to determine whether transcriptional patterns recapitulate those described in other species and to gain initial insight into the developmental timing of the GABA switch in swine.

We began by quantifying mRNA expression of *Slc12a2*, *Slc12a5*, and *Stk39* in the somatosensory cortex, hippocampus, and thalamus at embryonic day 36 (ED36), ED90, ED105 (sows gestate on average for 115-118 days) postnatal day 7 (“infant”), and postnatal day 30 (“toddler”). As shown in **Figure 1C**, *Slc12a2* expression remained relatively constant in the hippocampus and thalamus, while cortex levels exhibited a moderate upward trend over time. The level of *Slc12a2* transcript across biological replicates, regions, and ages was variable; this complexity may reflect underlying cellular heterogeneity as NKCC1 is expressed in both neurons and non-neuronal cells [29]. *Slc12a5* expression across all regions remained relatively stable. *Stk39* declined in hippocampus and thalamus between late gestation and infancy (2-way ANOVA main effect of age p=0.0007, main effect of region p=0.0008; post-hoc multiple t-test **p<0.01). Taken together, transcriptional analysis was not sufficient to resolve developmental changes in chloride cotransporters, though the drop in *Stk39* around birth might indicate a perinatal GABA switch. To gain a clearer understanding, we next turned to protein-level analyses.

#### Protein abundances and phosphorylation state for NKCC1, KCC2, and SPAK across development and region in swine

Representative western blots from randomly selected biological replicates in each group are shown in **Figure 1D**. Band densitometry was normalized to the total protein stain, as β-actin, β-III-Tubulin, and GAPDH were found to be poor loading controls across ages. Specifically, β-actin was more abundant in younger samples, and β-III-Tubulin and GAPDH were more abundant in older samples. Phosphorylated proteins were probed first, followed by total proteins, and in the case of NKCC1 and KCC2 exhibiting similar molecular weights, technical replicate full gels were run, and each sample was normalized to total protein in that gel. For each region, all ages and biological replicates were performed within the same experiment with a select replicate serving as an internal loading control for each membrane to account for any additional variance. In **Figure 1E**, total protein abundance of NKCC1, KCC2, and SPAK (left) and the phosphorylated ratios of those proteins are visualized (right). Strikingly, levels of NKCC1 remain relatively constant across ages and regions, while SPAK and KCC2 are significantly upregulated postnatally (KCC2: 2-way ANOVA main effect of age p=0.0002, main effect of region p=0.0043; post-hoc multiple t-test **p<0.01; SPAK: 2-way ANOVA main effect of age p<0.0001, main effect of region p-0.008; post-hoc multiple t-test *p<0.05,****p<0.0001). Phosphorylation ratios for SPAK and KCC2 all decline postnatally (pKCC2/KCC2: 2-way ANOVA main effect of age p=0.0015, main effect of region n.s.; post-hoc multiple t-test **p<0.01; pSPAK/SPAK: 2-way ANOVA main effect of age p<0.0001, main effect of region p=0.0104; post-hoc multiple t-test **p<0.01). Phosphorylation ratio for NKCC1 declines perinatally (pNKCC1/NKCC1: 2-way ANOVA main effect of age p=0.0012, main effect of region p=0.018; post-hoc multiple t-test p=0.0649). Taken together, the mRNA and protein data suggest that the developmental “GABA switch” in swine follows a largely perinatal trajectory and both PND7 and 30 piglets have undergone the GABA switch. This is supported by the general trend for increased total KCC2 and decreased pNKCC1 and pKCC2, pointing toward a maturation of inhibitory signaling. These findings are consistent with swine occupying a semi-altricial developmental niche and establishes a high-resolution baseline for these proteins in swine across brain regions. Given that our transcriptional data alone did not reliably reflect the abundance state of these transporters, this underscores the importance of assessing phosphorylation status and protein abundance when studying markers of GABAergic maturation.

With this framework in place, we investigated how these developmental differences might shape the brain’s response to injury. Rather than using a single-insult model like controlled cortical impact (CCI), which primarily captures focal mechanical injury, we employed a MuLMI TBI model designed to more closely replicate the complex pathophysiology observed in infants and toddlers with severe traumatic brain injury [27, 30]. This model has multiple pathoanatomic lesions (mass effect, subdural and subarachnoid hemorrhage) as well as insults (traumatic seizures and brief apnea and hypoventilation) under non-GABAergic sedation. The MuLMI model mimics components frequently observed in abusive head trauma [30]. By simulating this multifaceted injury environment, we aimed to better understand how developmental regulation of chloride transporters might contribute to age-specific patterns of tissue damage and excitability following trauma.

#### Creation of multi-factorial, severe TBI injuries and resulting age-specific pathology patterns

Implementing a MuLMI TBI model in a large animal is resource- and labor-intensive, but essential for studying complex pediatric brain injuries with high translational fidelity to human pathophysiology. **Figure 2A** illustrates the model, including the location of injuries, the placement of EEG electrodes for seizure monitoring, and the timeline of insults, injuries, and intensive care. Following GABA agonist weaning, the pathophysiology was allowed to evolve over a 24-hour period under intensive care conditions where the tissue damage develops beyond the area of the cortical impact. The right panel displays the ICU equipment used to sedate, intubate, and ventilate piglets. Examples of gross pathology (**Figure 2B)** and stylized maps of the damaged regions (yellow, **Figure 2C**) are provided, demonstrating that damage extends beyond the site of the cortical impact (arrow). The total damaged area was greater in piglets subjected to severe TBI injuries compared to sham injuries (Two-way ANOVA, main effect of treatment p = 0.02, **Figure 2D**). Furthermore, damage distribution exhibited age-dependent patterns (**Figure 2E**) where the percentage of hemisphere damaged due to TBI injuries was greater in the ipsilateral hemisphere compared to the contralateral hemisphere in toddlers compared to infants (Two-way ANOVA, main effect of treatment p=0.05, post-hoc t-test p = 0.0765). These findings highlight a clear age-specific vulnerability to this severe model of TBI and motivated our desire to understand why toddler piglets might have more tissue damage, possibly via upregulation of neuronal NKCC1.

#### TBI-associated shifts of transcription of cation chloride cotransporters were characteristic of reversion of the GABA switch in toddler piglets

We thus examined the effects of severe TBI on chloride transporter gene expression in piglets at different developmental stages, focusing on age- and region-specific transcriptional responses in the cortex. The cortex is where the differential patterns of injury are most obvious (**Figure 3A**). Quantitative PCR revealed *Slc12a2* (encoding NKCC1) expression was largely unchanged in cortex but exhibited a modest effect of age (**Figure 3B**; main effect of age p=0.0314). *Slc12a5* (encoding KCC2) was significantly downregulated in the cortex of injured toddler piglets (**Figure 3C**, main effect of age p=0.0039, main effect of treatment p=0.0636, age x treatment p = 0.0535). Stk39 (encoding SPAK), a kinase that regulates both NKCC1 and KCC2, was significantly upregulated of toddler piglets with TBI in cortex (**Figure 3D**, main effect of age p=0.0066, main effect of treatment p=0.0686, age x treatment p=0.0352).

Given that cortical transcriptional effects were most consistent across toddler piglets, we applied GeoMx spatial transcriptomics to cortical regions from toddler piglets with sham or severe TBI injuries. Regions of interest (ROIs) were chosen from superficial gray matter (teal) and gyral white matter (orange), as shown in the schematic (**Figure 3E**). We found that TBI-induced reductions in *Slc12a5* and increases in *Slc12a2* were more pronounced in white matter than gray matter, specifically in areas of structural damage (**Figure 3F**). These results suggest that white matter is particularly vulnerable to transcriptional dysregulation of chloride homeostasis following TBI in toddlers.

Taken together, these findings support the hypothesis that severe TBI disrupts chloride transporter mRNA expression in a developmentally and regionally specific manner. The apparent acute (24 hours after injury) shift toward excitatory GABA signaling in surviving toddler neurons may contribute to ongoing tissue vulnerability and highlights a potential window for targeting NKCC1 activity beyond the acute post-injury period.

#### TBI increases SPAK abundance in the cortex of infant piglets greater than toddler piglets, driven by acute response after injury and independent of hemisphere

Western blotting of cortical lysates (**Figures 3G, H**, quantified in **3I-J**), rostral and lateral to the area of cortical impact but still damaged, revealed that infant piglets with TBI exhibited relatively unchanged levels of NKCC1 and KCC2 due to TBI injuries, but SPAK protein levels greatly increased with TBI in infant piglets only (**Figure 3I**; Two-way ANOVA effect of treatment p=0.117; post-hoc multiple t-test infant sham v TBI p=0.0834; infant TBI v sham TBI p=0.0338). This effect became more apparent when considering the timeframe injuries were allowed to develop in piglets with TBI injuries (1 hours of injury versus 24 hours), wherein SPAK demonstrated a robust effect of age and time (Two-way ANOVA, time x age effect p=0.0007). This increase in SPAK was mirrored in abundance of TRKB, a kinase receptor known to influence NKCC1 and KCC2 expression [31] (Two-way ANOVA effect of time p=0.0067, time x age p=0.201). There was no effect of hemisphere across any proteins examined (**Figure 3K**). The significant upregulation of SPAK and TRKB in infant piglets with TBI, but not toddler piglets, is interesting for several reasons. 1) Our previous work identified that while both ages demonstrate vasogenic edema due to TBI, toddler piglets demonstrate a greater upregulation of matrix metalloproteinases and more vasogenic edema than infant piglets [28]. Upregulation of both SPAK and TRKB in infant piglets with TBI injuries might be driving evolving cytotoxic edema that does not result in vasogenic edema [13, 32]. 2) SPAK abundance can be promoted by NFkB signaling [33], which could be activated by either the subarachnoid hemorrhage or cortical impact-induced mechanical stress in this model. 3) Upregulation of SPAK and TRKB acutely after injury, but less so as the injuries progress, could profoundly impact cell-type phosphorylation states of the transporters which are missed at the level of resolution offered by whole tissue lysates, thus, requiring investigation of phosphorylation state of NKCC1 via immunohistochemistry.

#### Severe TBI in swine produces region- and cell-type specific alterations in pNKCC1 that are correlated with damage, subarachnoid hemorrhage, and seizure duration

At 24 hrs post-injury, TBI significantly increased pNKCC1 expression in non-parvalbumin-expressing cortical neurons (NeuN^+^/pNKCC1^+^/PV^-^) of infant piglets positively correlated with seizure duration and SAH area, while cortical PV^+^ populations and hippocampal neuronal populations showed no changes, indicating region- and cell-specific effects. We utilized immunohistochemistry to assess phosphorylated NKCC1 (pNKCC1) expression in NeuN co-labeled mature neurons in the cortex lateral or caudal to the cortical impact (**Figure 4A**), and each ROIs for each animal are presented in **Figure S1** where the pattern of damage assessed by H&E is highlighted in yellow (same cohort of piglets as summary data presented in Figure 2). Apoptotic neurons assessed by nuclear chromatin condensation via Hoechst/NeuN^+^ across cortical ROIs generally increased in toddlers but not infants (**Figure 4C-E**), consistent with our previous reports [6, 27]. In cortical ROI 2, the site caudo-medial to contusion (not the area of impact, but where the neuronal ischemia spreads beyond the contusion site in this MuLMI mode), TBI increased pNKCC1 expression in NeuN^+^ neurons, without a preference for PV^+^/NeuN^+^ interneurons (main effect of treatment p=0.0017, **Figure 4C**). When comparing pNKCC1 levels to % hemispheric damage, **visualized in Figure 4D, far right**, % damage shows a significant correlation with pNKCC1 in both ages (infant-Spearman r = 0.833; p=0.0154; toddler- Spearman r = 0.943; p=0.0167); however, only infant piglets demonstrated a positive (non-zero) correlation assessed by linear regression (p=0.0736). Multivariate analysis further confirmed the relationship between neuronal-pNKCC1 expression caudo-medial to the cortical impact site and spreading damage, SAH%, and seizure duration as positive predictors across both ages (**Figure 4F**; SAH x pNKCC1 Spearman r = 0.56; p=0.031; seizure duration x pNKCC1 Spearman r = 0.61, p=0.018). Further cell-specific analyses indicated that PV^+^ interneurons did not show enhanced pNKCC1 after TBI, suggesting that these inhibitory interneurons remain relatively unaffected. Expression of pNKCC1 in non-neuronal cells in the cortex was not different according to age or injury. In this early time post-injury (24 hrs), gliosis and re-building of the extracellular matrix are not expected. We have previously demonstrated an age-dependent increase in matrix metalloproteinase activity as early as 1 hr post-injury with extensive matrix metalloproteinase activity at 24 hrs, indicating active loss of the extracellular matrix at this time point [28].

#### Transcriptional, but not translational or post-translational changes to cation chloride cotransporters are observed in regions with minimal structural damage

In the hippocampus and thalamus, areas typically spared from hypoxic-ischemic damage in this model (**Figure 5A**), we again tested the effect of TBI injuries on age-dependent cation-chloride cotransporter dysregulation. In hippocampus, *Slc12a2* increased following TBI across both ages (**Figure 5B**, main effect of treatment p=0.0085). Conversely, *Slc12a5* was downregulated with age, but there was no TBI effect (**Figure 5C**, main effect of age p=0.0001). Stk39 exhibited a significant upregulation due to TBI injuries in the toddler piglets (**Figure 5D**, main effect of age p=0.0104, main effect of treatment p=0.0176, age x treatment p=0.0097). In thalamus, *Slc12a2* was significantly downregulated in the thalamus in both age groups (**Figure 5E**, main effect of treatment p=0.0002). *Slc12a5* exhibited a significant age and TBI effect, with infant piglets with TBI upregulating *Slc12a5* and toddler piglets with TBI downregulating *Slc12a5* (**Figure 5F**, main effect of age p=0.0038, main effect of treatment p=0.9538, age x treatment p=0.0293). Stk39 demonstrated a modest effect of TBI, but not by age (**Figure 5G**).

We observed minimal changes to NKCC1, KCC2, and SPAK in infant TBI piglets compared to toddler TBI piglets (**Figure 5H-J**). Therefore, regions not displaying overt injury can exhibit early changes in chloride transporter mRNA expression following TBI, but this may not be reflected in protein abundance or phosphorylation within this acute window. Further, structural damage might require additional time to manifest beyond the 24-hour paradigm of our study.

To be thorough in our investigation, we also employed immunohistochemistry across the dorso-ventral axis of the hippocampus (**Figure S2A-B**). Intriguingly, we observed pNKCC1 expression across postnatal ages in granule cells, neurons and glia of the polymorph layer, and neurons throughout the cornu ammonius, which seems to be different than expression typically reported in rodents. In all hippocampal regions, less than 5% of total neurons were apoptotic (**Figure S2C-F**). There was only an effect of injury on neuronal-pNKCC1 expression in the dorsal dentate gyrus (**Figure S2C**), but not in the dorsal CA2 or ventral DG or CA2 (**Figure S2D-F**). However, unlike in cortex, the effect on neuronal-pNKCC1 reflected a decrease in expression, rather than an increase. This likely reflects a downregulation of excitatory transduction proximal to the damage spreading distant from the cortical impact, perhaps as a compensatory mechanism.

Collectively, these data suggest that TBI-induced elevations in pNKCC1 predominantly occur in surviving cortical PV^-^ neurons of younger piglets, possibly contributing to a pro-excitatory state associated with seizures. These observations provide evidence that TBI influences cation chloride transporter expression and activation differently across brain regions and developmental stages. Younger piglets appear more prone to TBI-related shifts in these key proteins, raising the possibility that early therapeutic interventions targeting chloride cotransporters could be beneficial. An alternative possibility is that these changes were not detected in toddler piglets due to advanced tissue damage, but we attempted to resolve this with cell-specific expression in apoptotic vs. healthy cells.

#### Postnatal pNKCC1 downregulation in human hippocampus

To compare these developmental patterns with those observed in humans, we examined hippocampal samples from infants with known causes of unexpected death other than from TBI to determine developmental changes around the time of birth. The study design and tissue validation steps are outlined in **Figure 6A**. Previous transcriptional data from qPCR and microarray studies suggest that the GABA switch in humans occurs around birth in both the limbic system and neocortex [17, 34, 35]. Furthermore, evidence from slice electrophysiology, western blot analysis, and immunolabeling indicates hyperpolarizing GABA activity in the adult temporal lobe cortex and hippocampus [36, 37], which is altered in epilepsies [38].

We did not find a significant effect of age on the abundance of mRNA for *Slc12a2, Slc12a5*, and *Stk39* in relation to adjusted postconceptional age (PCA) [gestational age plus postnatal age]. As shown in **Figure 6B**, transcriptional levels of these genes remained relatively stable across postnatal ages, and no positive correlation emerged between the ratio of *Slc12a5: Slc12a2* and PCA (**Figure 6C**). These transcriptional results indicate that major transcriptional shifts in these transporters are minimal during the postnatal period in humans similar to development in piglets.

We then assessed protein abundances and phosphorylation states in these hippocampal samples (**Figure 6D**). Our study offers a more detailed resolution and analysis of the activity state (via phosphorylation) of these cotransporters during the immediate postnatal period in infants with normal brain development. Total protein abundances for NKCC1, KCC2, and SPAK all increased with PCA (**Figure 6E**; linear regression NKCC1 R^2^ = 0.7019, p=0.0025 non-zero; KCC2 linear regression R^2^ = 0.5247, p=0.0178 non-zero; SPAK linear regression R^2^ = 0.5953, p=0.0089 non-zero). An assessment of the ratio of active KCC2:NKCC1 (KCC2 / (pNKCC1/NKCC1) increased postnatally (linear regression R^2^=0.5610, p=0.0127 non-zero, **Figure 6F**), like swine. Postnatal KCC2 dominance aligns with the gradual shift toward inhibitory GABA signaling that characterizes brain maturation [9, 16, 17, 39, 40].

#### Upregulation of pNKCC1 expression is associated with tissue damage in autopsy cases of pediatric TBI

To bridge our findings from the piglet model to humans, we examined pNKCC1 expression in autopsy samples from non-TBI and TBI cases covering a range of ages (**Figure 7**). In non-TBI “control” human tissue, pNKCC1 was consistently present at low to moderate levels in neurons and glia across multiple regions (**Figure 7A**). In TBI cases, pNKCC1 expression varied according to injury severity and location. Regions with significant hypoxic-ischemic damage (the same type of tissue damage we model here), including the parietal lobe, corpus callosum, and hippocampus, showed increased pNKCC1 in glia and axons relative to controls (**Figure 7B, 7C**). The temporal lobe, especially the ependyma of the striatum terminalis, exhibited pronounced subependymal pNKCC1 upregulation in cases with severe white matter injuries.

## DISCUSSION

A central feature of normal brain development is the “GABA switch,” in which signaling becomes predominantly hyperpolarizing because of a change in the direction of chloride flux from outward (depolarizing) to inward (hyperpolarizing). This switch is correlated with shifts in NKCC1 and KCC2 expression, although a myriad of other factors affects the concentration of chloride such as impermeant anion moieties [41]. Our swine data show that this switch occurs around the perinatal period in a pattern more closely resembling semi-altricial species (including primates) than altricial rodents. In humans, we similarly observed a gradual postnatal decrease in NKCC1 phosphorylation aligned with a relative increase in potential inhibitory tone over time. Together, these findings suggest that swine provide a useful experimental bridge for modeling human neurodevelopment and the timing of the GABA switch.

The standard of care for seizures in infants and toddlers after TBI is attempting to stop seizures with GABA agonists, even though it is not known if seizures drive pathophysiology or are a symptom of TBI. Data on seizure incidence are confounded with TBI severity: worse brain injury results in more seizures. Furthermore, stopping seizures is difficult in infants with both levetiracetam and phenobarbital only working half the time [42]. We hypothesized that regions of tissue damage (i.e. cerebral cortex) would exhibit changes to the NKCC1 and KCC2 and regions without tissue damage (hippocampus and thalamus) would not, which clinically, would be expected to make stopping seizures even more difficult. We did find upregulation of neuronal pNKCC1 in the cortex following TBI injuries as predicted.

We further hypothesized that toddler piglets with more pervasive, hemispheric spreading injury with more pronounced vasogenic edema would exhibit upregulated NKCC1 compared to infant piglets with patchy, less tissue damage. However, we found the opposite to be true where greater upregulation of SPAK in response to injury was not in the age with greater tissue damage. Younger animals displayed a more pronounced TBI-induced upregulation of SPAK, a kinase that promotes NKCC1 phosphorylation and activity. This shift toward elevated NKCC1 function can increase neuronal excitability and seizure risk—clinical challenges frequently observed in pediatric TBI. This is perhaps because apoptotic (dead) neurons, possibly due to cytotoxic edema, which are abundant in the injured toddler piglets, are non-functional and will not be affected by changes to the Cl^-^ milieu, whereas surviving neurons might exhibit ongoing dysfunction. The correlation between pNKCC1 counts and seizure duration and hemorrhage in piglets further reinforces the notion that disrupted chloride regulation might contribute to post-traumatic neuronal excitation. Notably, these TBI-induced changes were not uniform across cell types or regions: parvalbumin-positive cortical interneurons were spared, and hippocampal neurons remained relatively unaffected by these specific injuries. Spatial transcriptomics indicated that white matter alterations in NKCC1 and KCC2 expression likely involve glial cells and support matrix, influencing extracellular ion concentration [41] and potentially contributing to edema and barrier dysfunction. Indeed, we previously demonstrated that toddlers, which have the most damage, also have ongoing upregulation in abundance and activity of matrix metalloproteinase, whereas matrix metalloproteinases are either not increased or even downregulated in infants, who have less damage [28]. This work supports the hypothesis that chloride transporters alone do not control intraneuronal chloride concentrations, especially after pediatric TBI [41, 43]. Human autopsy data corroborated these animal findings. In TBI cases, we saw similar heterogeneous region- and cell-type-specific shifts in pNKCC1, particularly in damaged white matter tracts.

An alternative hypothesis is that reversion back to NKCC1, a biomarker for increased intraneuronal chloride, is protective in PND7 piglets, who demonstrate less tissue damage, and perhaps at the expense of future hyper-excitability and development of post-traumatic epilepsy. Indeed, the only way to determine cause vs. effect in the developmental differences in response to injury is to provide therapies that would reduce MMP upregulation such as matrix metalloproteinase inhibitors and/or NKCC1 an antagonist such as bumetanide. Another level of complexity is that NKCC1 expression in non-neuronal cells have been shown to mediate cell death and vasogenic edema in models of ischemia (reviewed by [44]). Here, the differential neuronal-specific NKCC1 increase after severe TBI was not detectable via Western blots of mixed cell lysates.

Taken together, after TBI, toddler piglets display vasogenic edema likely from MMP upregulation, while infant piglets might instead experience acute cytotoxic edema with rapid, clean cell death without leading to vasogenic edema, resulting in ongoing neuronal dysfunction in surviving neurons, which might make them unique candidates for therapeutic intervention [45]. This postulate is supported by Level III clinical evidence and recent age-dependent mechanisms of cerebral edema in small-diameter blood vessels following severe TBI in an autopsy case-series investigation [46]. Given the robust age- and treatment-effects of TBI injuries on SPAK abundance, we propose that treatment of our infant piglets with a SPAK inhibitor immediately following TBI might offer protective and antiepileptic effects [47, 48]. Indeed, as pathophysiology is age-specific, successful therapeutics might also need to be age-specific after severe TBI according to developmental stage: targeting NKCC1 or SPAK in infants and targeting MMP’s in toddlers.

### Limitations

Our study has several limitations that should be acknowledged. Firstly, we were not powered to examine the distinct effect of sex as a covariate on NKCC1 and KCC2 levels in the developing swine brain due to protocol limitations, which may influence the observed outcomes [49]. Additionally, we did not assess the levels of brain-derived neurotrophic factor (BDNF) or insulin-like growth factor (IGF), both of which are known to promote the GABA switch. Other intracellular regulators such as protein kinase C (PKC), Src kinases, and with no-lysine (WNK) kinases, the latter of which are chloride-sensitive and may play a role in the switch, were also not considered in this study [48, 50]. Furthermore, we did not investigate the potential contributions of maternal estrogen or taurine as positive sensory regulators of the GABA switch. Finally, and perhaps most importantly, we did not perform electrophysiology or chloride ion imaging either *in vivo* or in organotypic slice, thus, our observations are limited to when the switch is *<u>likely</u>* occurring, however, we cannot say with certainty when GABA switches from excitatory to inhibitory in the developing swine brain.

Our study, while leveraging a translationally relevant swine model and human autopsy samples, cannot fully capture the clinical complexity and heterogeneity of pediatric TBI. The standardized injury approach in pigs, though informative, may not reflect the multifaceted trauma mechanisms, variable interventions, and long recovery trajectories seen in patients. These factors represent important avenues for future research to better understand the mechanisms underlying the changes in excitation and inhibition in the developing swine brain and in the context of TBI. Donnan effects with the loss of the extracellular matrix also likely affect the amount of extraneuronal and intraneuronal chloride concentrations post-TBI as demonstrated in rodent hippocampal explants [41, 43]. Indeed, in our own severe TBI model, we have described an age-dependent upregulation of matrix metalloproteinases where there was an ongoing upregulation in toddlers with worse injury, compared to a resolution in infants with lower injury burden scaled to brain volume [28]. The human autopsy samples, while invaluable, represent end-stage conditions and lack information on the temporal progression of TBI pathology. Further, it is unclear to what extent non-traumatic pediatric autopsy samples are representative of a physiologically normal pediatric population.

We observed notable variability in transcript-level measurements, likely due to biological heterogeneity and technical factors such as RNA quality, sampling consistency, and cellular composition of lysates. Though immunoblotting analyses were particularly informative, phosphorylation state was less informative in swine. It is important to note that our study did not evaluate additional post-translational modifications, such as glycosylation, though these are certainly indicated in the activity of these transporters. Finally, while we identified potential mechanistic links between chloride transporter dysregulation, seizures, and injury severity, further studies are needed to establish causal relationships and to explore how interventions that target these pathways might improve long-term outcomes in pediatric TBI.

### Conclusions and future directions

By integrating findings from a swine model, whose brain development and structure closely parallel those of humans, with postmortem human tissue analyses, our work supports the translational value of targeting chloride regulation to improve TBI outcomes, especially in pediatric populations [13, 51, 52]. Pharmacologically modulating NKCC1 or KCC2, potentially with agents like a SPAK inhibitor, could help restore chloride homeostasis and reduce hyperexcitability and swelling/tissue shifts after pediatric TBI. Such treatments would need to consider the patient’s developmental stage and the specific brain regions affected. Early intervention, when the brain is most vulnerable to ionic changes that might exacerbate tissue damage, may be especially beneficial. Our findings emphasize that TBI’s impact on the developing brain cannot be fully understood by focusing solely on neuronal excitability or a single brain area. Instead, we must consider a dynamic interplay among developmentally regulated ion transporters, glial function, neuroinflammatory processes, and structural integrity. Future work should address 1) how these factors interact over longer recovery periods, potentially via *in vivo* monitoring of neuronal chloride [53], 2) explore targeted therapies that normalize chloride concentrations, and 3) consider how interventions might influence long-term neurodevelopmental outcomes and post-traumatic sequelae such as epilepsy.

## Acknowledgements

This work is supported by NIH NICHD K01HD083759 and R01 HD099397 (BCB), by the Pappendick Family Translational Research Program (MKL), the Hydrocephalus Association, and R01 HD090064 (RH). AH was supported by T32NS007473 and F32NS134588 training fellowships from the National Institute for Neurological Disorders and Stroke. YC was supported by the Howard Hughes Medical Institute through the James J. Gilliam Fellowships for Advanced Study program. PO was supported by a Mass General Summer Research Trainee Program (SRTP). We thank the Boston Children’s Hospital IDDRC Cellular Imaging Core supported by S10OD016453. Spatial transcriptomics was facilitated by the Comparative Pathology and Genomics Shared Resource core facility at The Cummings School of Veterinary Medicine at Tufts University. We thank the past and present team of undergraduate interns in the Costine-Bartell laboratory who assisted with the surgeries and 30 hours of intensive care to create the brain injury model and tissue collection as well as the Knight Surgical Research Lab for assistance with anesthesia to create the brain injuries. We are especially grateful for the fetal piglet tissue obtained from Dr. Ryann Fame (Stanford University Department of Neurosurgery), Dr. Mercedes Paredes (University of California, San Francisco (UCSF) Department of Neurology), and Dr. Misty Williams-Fritze (CBSET) and postnatal piglet tissue obtained from Dr. Corinna Beale (Center for Comparative Medicine, MGH). We thank Dr. Cameron Sadegh (UC Davis Department of Neurosurgery) and members of the Lehtinen Laboratory for assistance with prenatal swine brain dissections. We are deeply grateful to participating families for their courage and generosity in consenting to the use of their children in research. We also thank the clinicians, medical examiners, and pathologists who helped in the accrual of cases, including Drs. Gary Dale, Jeffrey Goldstein, Steven Campman, Catherine Stoos, and Colin Smith. We thank Elisabeth Haas and Richard Goldstein for the processing and analysis of infant cases. We thank Jane’s Trust Initiative, the NIH NeuroBioBank, and the Edinburgh Brain Bank. We thank Kevin Staley for careful reading and suggestions on the research article.

## Author contributions

AH: Conceptualization, investigation, methodology, formal analysis, writing-original draft, visualization; YC: Investigation, methodology, formal analysis, writing-original draft; PO: Investigation, methodology, formal analysis, visualization; BB: Investigation, methodology, formal analysis, visualization; AD: Investigation, methodology; DM: Investigation, formal analysis, writing-revising; TS: Investigation, methodology; RH: Investigation, methodology, formal analysis, writing-revising, funding acquisition; MKL: Writing-revising, funding acquisition; BCB: Conceptualization, investigation, methodology, visualization, writing-original draft, project administration, funding acquisition

---

**Supplemental Figure 1:**
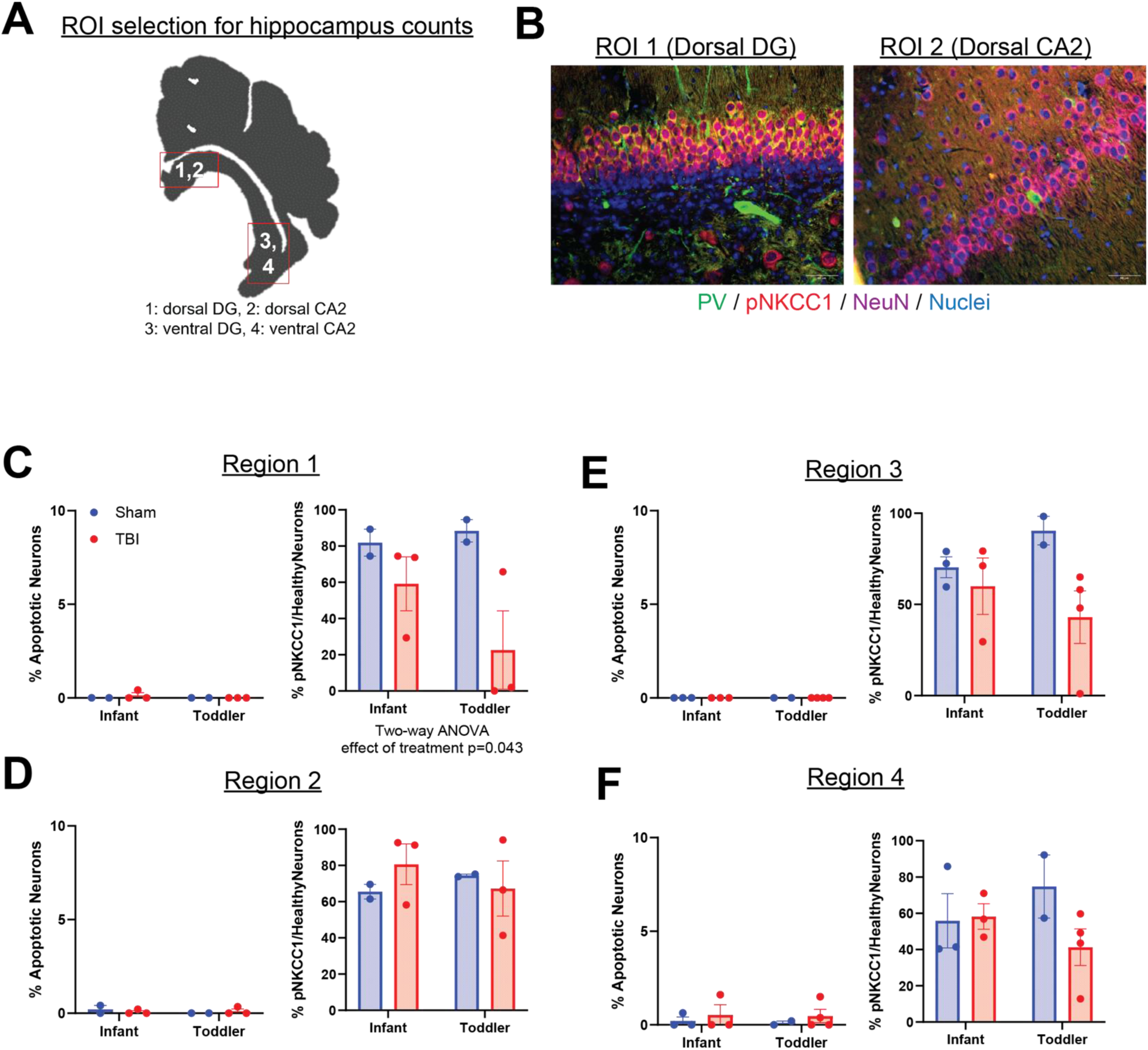
Damage maps for each animal and region used for cell counts in **Figures 4 and S2**. Yellow map shows damage visible on H&E. Red box indicates approximate ROI used for counts. ***A***. ROIs for Cortex 1 ***B*** ROIs for Cortex 2 and 3. ***C,*** ROIs for hippocampus counts.

**Figure S2:**
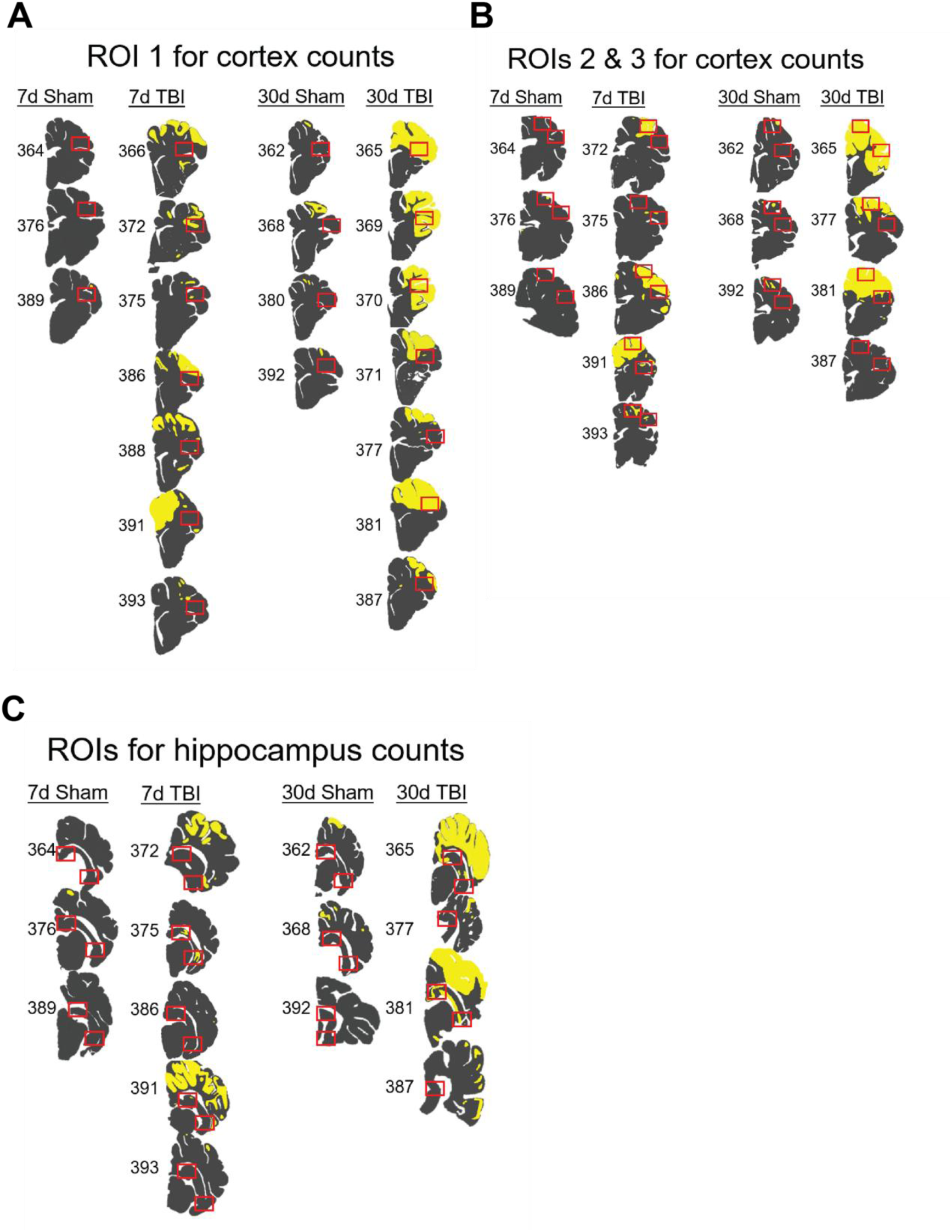
The expression of neuronal pNKCC1 expression in the hippocampus of swine was not affected by age or injury. Schematic of ROIs for analysis, ***A***. Immunohistochemical staining for pNKCC1 (red), parvalbumin (green), NeuN (magenta), and nuclei (white), ***B***. Cell counts expressed as percentages per ROI, with ROI 1 in ***C***, ROI 2 in ***D***, ROI 3 in ***E,*** and ROI 4 in ***F***. In all cases, the **left** dot plot presents % of neurons that were apoptotic, and the **right** dot plot presents the % of healthy neurons expressing pNKCC1. Two-way ANOVA with post-hoc multiple t tests employed for all comparisons and significant findings for main effect of age or treatment reported in graphs.

