## Supplemental Figures for "Temporal and cell-type specific SPAK-NKCC1 disruption following severe TBI in the developing gyrencephalic brain"

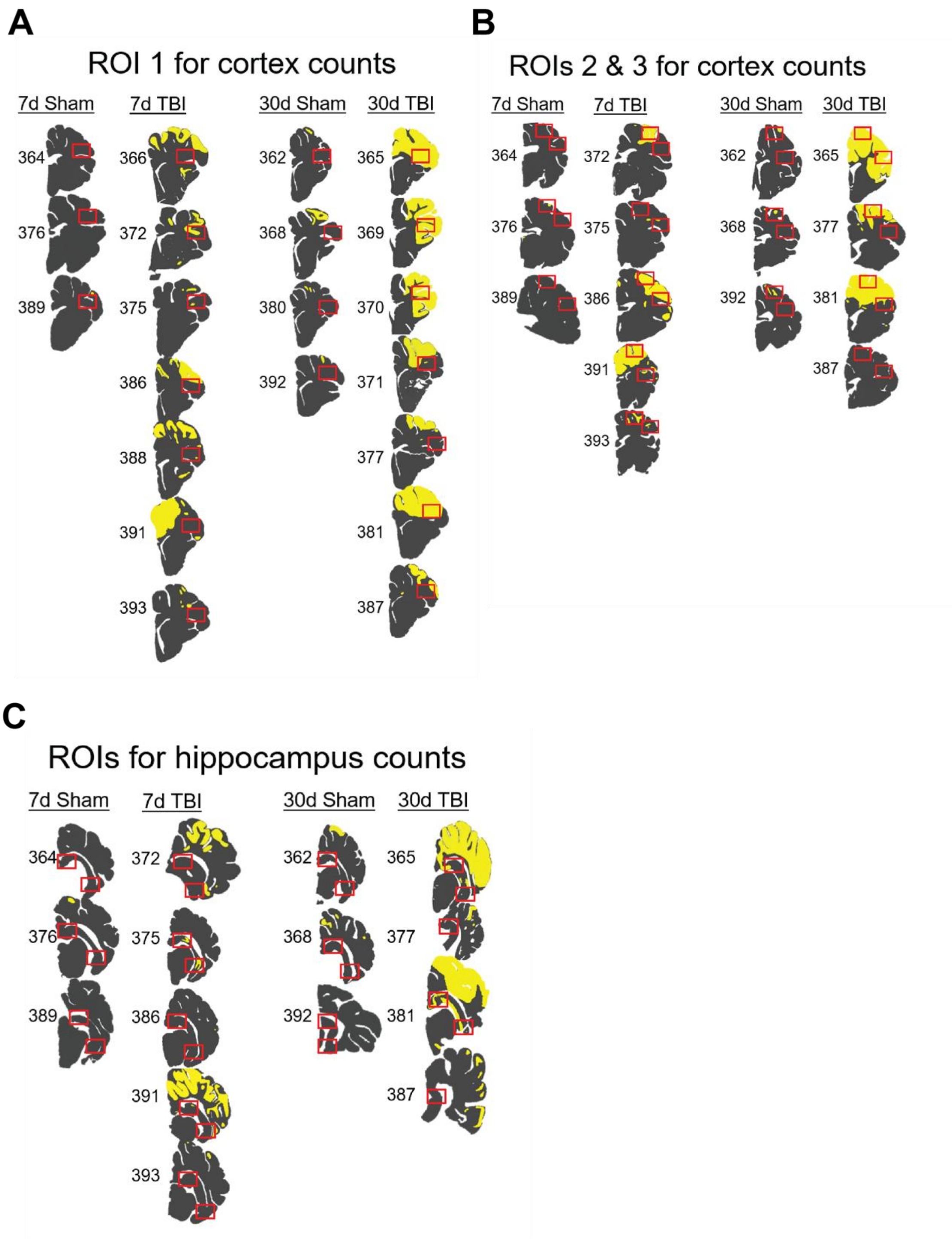


**Supplemental Figure 1:**

Damage maps for each animal and region used for cell counts in **Figures 4 and S2**. Yellow map shows damage visible on H&E. Red box indicates approximate ROI used for counts. ***A***. ROIs for Cortex 1 ***B*** ROIs for Cortex 2 and 3. ***C,*** ROIs for hippocampus counts.

**Figure S2:**

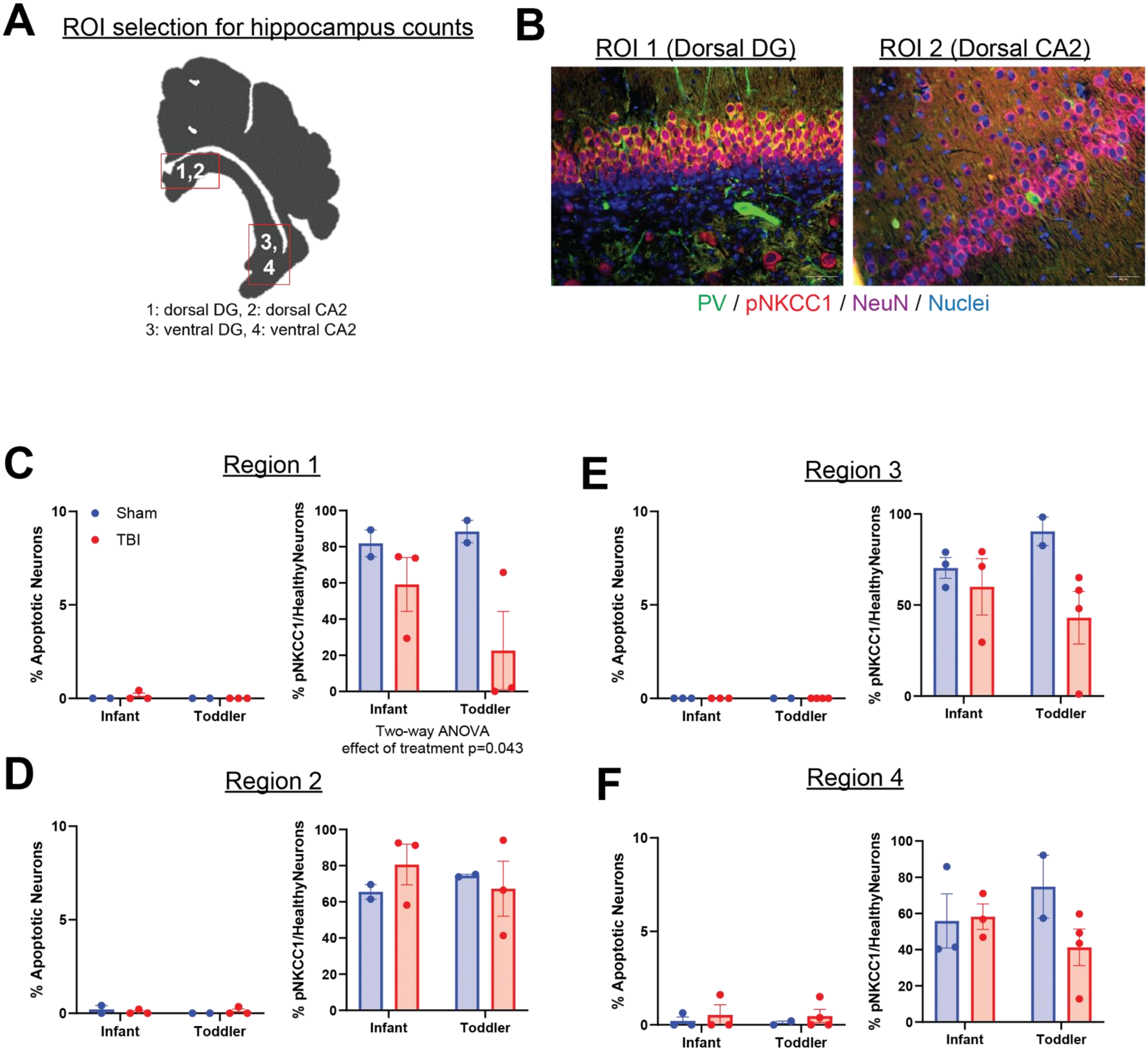


**Figure S2: The expression of neuronal pNKCC1 expression in the hippocampus of swine was not affected by age or injury.**

Schematic of ROIs for analysis, ***A***. Immunohistochemical staining for pNKCC1 (red), parvalbumin (green), NeuN (magenta), and nuclei (white), ***B***. Cell counts expressed as percentages per ROI, with ROI 1 in ***C***, ROI 2 in ***D***, ROI 3 in ***E,*** and ROI 4 in ***F***. In all cases, the **left** dot plot presents % of neurons that were apoptotic, and the **right** dot plot presents the % of healthy neurons expressing pNKCC1. Two-way ANOVA with post-hoc multiple t tests employed for all comparisons and significant findings for main effect of age or treatment reported in graphs.
